# BPEA integrates pleiotropy and epistasis to guide allele pyramiding for multi-trait crop improvement

**DOI:** 10.64898/2026.11.19.767812

**Authors:** Xiaodong Li, Yuhong Wang, Boyu Meng, Wei Chang, Jie Xiang, Zhixuan Qin, Jiaming Song, Kun Lu

**Author notes:** Correspondence: Jiaming Song. College of Agronomy and Biotechnology, Southwest University, Chongqing 400715, China.; Kun Lu. College of Agronomy and Biotechnology, Southwest University, Chongqing 400715, China. The author responsible for distribution of materials integral to the findings presented in this article in accordance with the policy described in the Instructions for Authors (https://academic.oup.com/plcell/pages/General-Instructions) is: Kun Lu.

## Abstract

Dissecting the genetic basis of complex traits and pyramiding favorable alleles are central to crop breeding. However, jointly evaluating pleiotropy and epistasis to identify allele combinations that meet multiple breeding objectives remains challenging. Here, we present BPEA (Bayesian Pleiotropy and Epistasis Analysis), a Bayesian framework integrating LD-guided QTL interval identification, joint genetic inference and allele-combination assessment. BPEA uses the joint posterior to estimate the posterior probability of simultaneous improvement (PPSI) between allele combinations and ranks them by mean PPSI. In simulations, BPEA improved interval identification and epistatic interaction attribution relative to comparator methods and maintained high precision in shared-signal detection. When epistasis was present, it reduced mean combination-selection regret by 68.4–86.3% relative to a model without interaction terms. Applications to three rapeseed populations identified shared genetic signals and candidate epistatic interactions associated with seed-quality traits. BPEA also prioritized allele combinations for simultaneous reductions in erucic acid and glucosinolate contents and identified complementary parents for cross design. BPEA thus provides a framework for genetic dissection and genomics-guided crop improvement.

## Introduction

Complex crop traits, including yield, quality and adaptation, are shaped by the combined effects of multiple loci, environmental conditions and their interactions. Pleiotropy and epistasis are fundamental features of the genetic architecture of quantitative traits (Mackay and Anholt, 2024). Pleiotropy contributes to this genetic complexity by allowing a single locus to influence multiple traits. For example, natural variation in rapeseed *ARF18* affects both silique length and seed weight (Liu et al., 2015). Epistasis makes the effect of one locus depend on the genotype at another. In tomato, negative epistasis can cause excessive inflorescence branching and reduced fertility (Soyk et al., 2017). Dissecting the genetic basis of complex crop traits therefore requires identifying trait-associated loci and characterizing their pleiotropic effects and epistatic interactions.

Population resequencing and extensive phenotyping have made genome-wide association studies (GWAS) an important approach for identifying genetic variants significantly associated with crop traits (Hu et al., 2022). Mixed models account for relatedness among individuals and reduce confounding by genetic background (Zhou and Stephens, 2012). However, single-marker association tests identify associated markers but do not directly delineate quantitative trait locus (QTL) intervals. In crop GWAS, candidate intervals have been defined using fixed windows around associated markers or local linkage disequilibrium (LD). A rapeseed study, for example, selected genes within local LD blocks and used windows extending 100 kb on either side of associated markers when no block could be identified (Knoch et al., 2020). The p-clump method implemented in PLINK (Chang et al., 2015) has also been used in crop GWAS to group markers based on association significance and LD (Qi et al., 2025; Wang et al., 2025), but LD with a lead marker alone does not establish whether the resulting intervals contain distinct LD segments. Because interval boundaries determine candidate-gene sets and the variants included in joint analysis, interval delineation should account for local LD structure and include an assessment of boundary stability.

Genomics-guided pyramiding of favorable alleles has contributed to crop improvement (Gu et al., 2023; Yao et al., 2026). Using candidate QTL intervals to guide allele pyramiding requires understanding both shared genetic effects across traits and epistasis between loci. coloc evaluates evidence for shared associations (Giambartolomei et al., 2014), while its integration with SuSiE accommodates multiple association signals within a region (Wang et al., 2020; Wallace, 2021). FaST-LMM tests epistasis between loci (Lippert et al., 2013). Multiple-trait multiple-interval mapping and mvMAPIT also use information across traits to investigate pleiotropy or epistasis (Da Costa e Silva et al., 2012; Stamp et al., 2023). However, detecting shared genetic signals or significant interactions alone does not quantify the probability that specific allele combinations will improve multiple traits simultaneously. A new analytical strategy is therefore needed to jointly infer shared main effects and epistatic interactions and propagate uncertainty in genetic configurations and effects into allele-combination assessment.

Here, we develop BPEA (Bayesian Pleiotropy and Epistasis Analysis), a framework that integrates LD-guided QTL interval identification, joint Bayesian inference and allele-combination assessment. The bivariate model jointly infers shared genetic signals and candidate epistatic interactions and uses the same posterior to estimate the posterior probability of simultaneous improvement (PPSI) between allele combinations under specified genetic backgrounds. We evaluate interval identification using independently simulated genotype datasets and assess joint inference and combination selection using phenotype simulations based on real genotypes from three rapeseed populations. We then construct a QTL atlas across populations and trait-environment combinations, jointly analyse pairs of seed-quality traits, and rank allele combinations for simultaneous reductions in erucic acid and glucosinolate contents. By extending joint genetic inference to combination assessment and parent selection, BPEA provides a quantitative framework for pyramiding favorable alleles towards multiple breeding objectives.

## Materials and methods

### Data collection and quality control

We collected 17.45 Tb of public rapeseed resequencing data from NCBI BioProjects PRJNA476657, PRJNA358784 and PRJNA416679, designated Bna1007, Bna588 and Bna418, respectively. The ZS11 genome and annotation (Song et al., 2020) provided the reference for variant calling. Variants were retained at minor allele frequency (MAF) ≥ 0.05 and missingness ≤ 0.2. The Bna418, Bna588 and Bna1007 genotype datasets used here contained 418, 655 and 1,005 accessions and 2,335,178, 6,070,444 and 4,928,085 variants, respectively. Phenotypes covered phenology, plant architecture, morphology, yield components, seed quality, and stress and disease responses with 138 trait–environment combinations, comprising 89, 39 and 10 in the respective populations. Phenotypes were centred and scaled by their standard deviations before joint modeling.

### Genome-wide association analysis

Single-marker mixed-model association analysis used GEMMA 0.98.5 (Zhou et al., 2012), with marker dosage as a fixed effect, kinship as the covariance basis for polygenic effects, and an intercept. Analyses used -lmm 4, with Wald-test *P* values defining candidate association markers. A standardized relatedness matrix was generated from genome-wide genotypes using -gk 2 and supplied through -k. GEC estimated the effective number of independent tests, *N* (Li et al., 2012), and calculated the significant threshold. Finally, thresholds were 1.09 x 10□□, 3.97 x 10□ □ and 3.96 x 10□ □ for Bna418, Bna588 and Bna1007, respectively.

### LD-guided intervals and candidate-gene annotation

BPEA 0.1.0 was used to identify candidate QTL intervals. Unassigned significantly associated markers were selected in ascending order of GWAS *P* value. Unassigned markers within a 500-kb window on the same chromosome and with LD *r*^2^ > 0.2 to the representative marker formed an initial cluster. The secondary *P*-value threshold was 1, allowing linked markers without suggestive association to contribute to interval boundaries. Endpoints were the minimum and maximum member coordinates. By default, *r*^2^ is the squared Pearson correlation of allele dosages. An optional haplotype-LD mode estimates two-locus haplotype frequencies by maximum likelihood and calculates *r*^2^.

A binary support sequence records whether each marker meets the LD threshold relative to the representative marker. Internal boundaries are proposed in consecutive unsupported segments, requiring at least two supported markers and one entry marker on either side. Let *l*(*k,n*) = *k* log(*k*/*n*) + (*n*−*k*) log(1−*k*/*n*), where *k* is the number of supported markers and *n* is the number observed. The segmentation score is

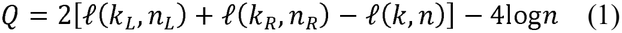

where L and R denote the left and right segments. The highest-scoring split with Q > 0 is retained and processed recursively; 4 log *n* penalizes excessive segmentation. All flanking markers within the original interval are then used to calculate between-flank and within-flank mean LD, excluding markers inside the gap. Boundary stability is assessed by

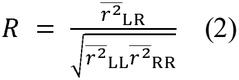

The numerator is mean *r*^2^ across marker pairs spanning the boundary; the denominator is the geometric mean of within-flank mean *r*^2^. Thus, *R* < 1 indicates weaker linkage across the boundary than within its flanks. Holding candidate boundaries fixed, individuals are resampled with replacement 199 times. A boundary is retained when both the observed *R* and the 95^th^ percentile of bootstrap *R* are below 1. Each subinterval must contain at least one entry marker. Fully nested intervals are absorbed into their containing intervals.

For the population QTL atlas, physically overlapping intervals were merged within each trait and chromosome. Neighboring intervals were also merged when separated by ≤ 500 kb and every cross-interval representative-marker pair had *r*^2^ > 0.2. Genes intersecting original intervals were identified from the gene annotation as candidates. Cross-population coverage similarity for the same trait was the Jaccard index, calculated as intersection length divided by union length.

### Interval simulation and evaluation

We generated 600 independent genotype datasets, each with 600 individuals and 600 markers spaced 10 kb apart across approximately 6 Mb. Six LD layouts, each replicated 100 times, varied region length, internal LD heterogeneity and linkage between neighboring regions. Each dataset generated phenotypes at PVE 8% and 16%, giving 1,200 non-null phenotypes with 900 target intervals per PVE level, plus 600 null phenotypes.

Genotypes coded 0/1/2 were generated from two binary haplotypes. Background allele frequencies followed Uniform (0.15, 0.50), ancestral regional template frequencies followed Uniform (0.35, 0.65), and marker-flip probabilities in ordinary regions followed Uniform (0.06, 0.12); heterogeneous regions used segment-specific flip probabilities. Individual homozygosis probabilities followed Uniform (0.90, 1.00). Standardized dosages at designated effect loci were summed with equal positive effects, and the genetic value *g* was scaled to unit variance. Phenotypes were generated as 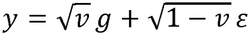, with standard normal ε and *v* = 0.08 or 0.16. Prespecified physical intervals defined the truth; adjacent low-LD subregions containing separate effect loci were distinct targets.

Simulated GWAS used linear regression with an intercept and two-sided *t*-tests, with a significant threshold of 0.05/600. Comparators were p-clump in PLINK 2 a6.5 (Chang et al., 2015) and fixed windows. p-clump settings were clump-p1 = 0.05/600, clump-p2 = 1, clump-r2 = 0.2 and clump-kb = 500; dosage correlation used clump-unphased. Reported intervals covered representative and member markers. Fixed windows selected representatives by *P* value and extended ±50, ±100, ±250 or ±500 kb. LD sensitivity analyses compared thresholds of 0.2, 0.4, 0.6 and 0.8 and dosage-correlation versus haplotype-based LD.

One-to-one matching required IoU ≥ 0.5, first maximizing the number of matches and then total IoU. True positives (TP), false positives (FP) and false negatives (FN) were matched intervals, unmatched reported intervals and unmatched targets. Recall was ΣTP/(ΣTP+ΣFN), and precision was ΣTP/(ΣTP+ΣFP). The interval-IoU score was the maximum total IoU without a matching threshold, divided by the larger of the reported and target interval counts, then averaged across phenotypes. Empirical interval FDR was mean FP/max (reported count,1) across phenotypes. Genotype-level paired bootstrap resampling with 10,000 replicates yielded 95% confidence intervals. Null evaluation used the proportion of phenotypes with at least one reported interval. Genotype and LD features were additionally compared with 80 regions spanning 19 chromosomes in 923 Bna1007 accessions, using 512 markers per region.

### Joint Bayesian analysis of pleiotropy and epistasis

For the *n* x 2 phenotype matrix *Y*, the joint mixed model is

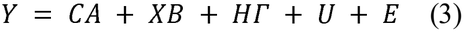

*C* is the covariate design matrix; *X* contains allele dosages, originally coded 0/1/2 and centred on sample means; *H* contains products of two centred dosages, representing additive-by-additive interactions. *A, B* and *Γ* are the corresponding coefficients, *U* is polygenic background and *E* is residual error. With observations ordered by individual, total covariance is

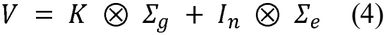

*K* is the kinship matrix, and Σ_*g*_ and Σ_*e*_ are genetic-background and residual trait covariances. These are estimated by restricted maximum likelihood (REML) under the null model and held fixed for configuration comparisons.

Genetic configuration *M* specifies main-effect sets and interaction assignments for both traits. Main and interaction coefficients have independent *N* (0,0.3^2^) priors on the input phenotype scale. Per-variant prior probabilities for trait-A-only, trait-B-only, shared and null states are 10□□, 10□□, 10□□ and 1−2 x 10□□−10□□. Prior masses for no interaction, A-only, B-only and both-trait interactions are 0.90, 0.04, 0.04 and 0.02; non-null masses are divided equally among candidate pairs. Configuration priors multiply variant-state and interaction-state probabilities and normalize over the allowed set □. Each configuration includes at most two main-effect loci per trait, three in their union and one active interaction pair.

Covariate coefficients have a common flat prior across configurations. After covariance whitening, covariate effects are removed by orthogonal projection, yielding design matrix *D* and phenotype vector *y*\*. *W* is the covariance of the independent normal effect prior. For configuration *M*, define

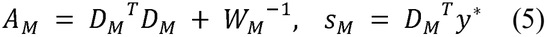

Integrating the normal effect coefficients gives the log Bayes factor relative to the null model:

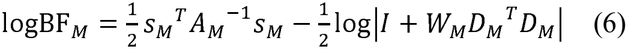

All posterior probabilities are conditional on the candidate set and estimated covariance. Configuration probabilities are

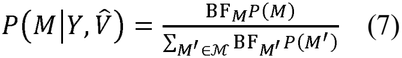

For main-effect sets *S*_*A*_ and *S*_*B*_ in configuration *M*, the regional sharing probability sums posterior mass over configurations containing at least one common locus:

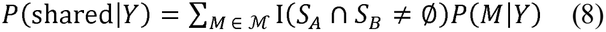

Let *E*_*M*_ contain the interaction effects in M. The PIP for pair (a,b) acting on trait t is

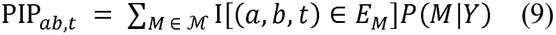

I(·) denotes the indicator function. Enumerable models are summed exactly; otherwise, four Markov chain Monte Carlo (MCMC) chains sample discrete configurations, each with 500 warm-up and 2,000 retained iterations. Non-constant monitored traces require split R-hat ≤ 1.05 and Monte Carlo standard error ≤ 0.02. Failing runs are extended to 2,000 warm-up and 8,000 retained iterations. Sharing probabilities and interaction PIPs use a support threshold of 0.95.

### Joint-model simulations and method comparisons

The 864 bivariate simulations used real genotypes from Bna418, Bna588 and Bna1007, with sample sizes of 403, 655 and 1,005. Six regions per population each generated 12 phenotype templates, crossed with presence or absence of sharing and epistasis to give 288 cases per population. Half of the sharing-positive templates contained one shared main effect; the remainder also contained a trait-B-specific effect. Sharing-negative templates used distinct main-effect loci.

Relative to a unit baseline target variance, main effects contributed 0.08, polygenic background contributed 0.30, and residual trait correlation was 0.743441. Interaction-positive settings added a variance component of 0.08 without rescaling total variance. The four settings for a template shared background and residual draws. Each case nominated four genotype-based candidate pairs with MAF ≥ 0.05, inter-locus *r*^2^ ≤ 0.2 and estimable interaction effects. The true interaction rotated among these pairs and affected trait A, trait B or both.

Sharing comparators were coloc-ABF and coloc-SuSiE (Giambartolomei et al., 2014; Wang et al., 2020; Wallace, 2021), using GWAS statistics for identical candidate variants and priors *p*_*1*_ = *p*_*2*_ = 10□□ and *p*_*12*_ = 10□□. Interaction comparators were pairwise P3D-LMM, joint P3D-LMM and FaST-LMM 0.6.13 (Lippert et al., 2013; Zhang et al., 2010). Pairwise P3D included two main effects and their product; joint P3D included all four candidate products and their main effects. Both used null-model REML variance components. FaST-LMM compared main-effect and interaction models by likelihood-ratio tests. The eight tests per case, covering four pairs and two traits, underwent Benjamini–Hochberg correction (Benjamini and Hochberg, 1995), with *q* ≤ 0.05 defining support.

Sharing was evaluated per region and trait pair, giving 288 units and 144 positives per population. Interactions were evaluated per exact pair and target trait, giving 2,304 units and 192 positives per population. TP, FP and FN were pooled to calculate recall, precision and empirical FDR = FP/(TP+FP). Confidence intervals used 2,000 stratified paired bootstrap replicates, resampling regions and then templates within regions. Single-threaded runtime on an Intel Xeon Gold 6248R included method-specific association calculations, covariance estimation and inference. BPEA timing covered both sharing and interaction outputs.

Ablations used the same 864 cases, candidate sets, phenotypes, main-effect configuration priors and 0.95 posterior threshold. No trait covariance set off-diagonal elements of genetic and residual covariance matrices to zero, retaining trait variances, kinship and sharing priors. No interactions omitted interaction terms. Pairwise interactions retained all candidate main effects and fitted each of the four pairs separately, restricting and renormalizing the original prior to no interaction or the current pair while preserving prior odds. The additive-only model evaluated sharing; pairwise models evaluated interactions. We also reported F_1_ = 2TP/(2TP+FP+FN). Each model used four chains with 500 warm-up and 2,000 retained iterations, extended to 2,000 and 8,000 when monitored split R-hat exceeded 1.05 or Monte Carlo standard error exceeded 0.02. Differences used the same stratified paired bootstrap.

For combination-selection validation, 64 observed genotype profiles were sampled using genotype information alone for each region–template pair and held constant across its four simulation settings. The full training sample was retained. BPEA, No interactions and No trait covariance each used 20,000 posterior configuration-and-effect draws to rank profiles by their mean probability of jointly lowering both traits relative to the other 63 profiles. True scores were the fractions of those profiles strictly improved in both traits under the noiseless simulated local main and interaction effects. Selection regret was the mean true score of the optimal top 20% minus that of the model-selected top 20%, with fractional weighting of tied scores at the selection boundary. Paired comparisons included 863 cases after excluding one case with unresolved sampling in No interactions; 245 cases with identical true scores were retained. Mean regret and model differences used 2,000 paired region–template bootstrap resamples, conditional on the fitted models and sampled profiles.

### Joint analysis of real populations

BPEA analyzed 71 seed-quality trait pairs across the three populations using 3,402 within-region or cross-region models, and plant architecture and yield components using 1,597 models. Within-region main-effect candidates were QTL variants with MAF ≥ 0.01 and missingness ≤ 0.05, recalculated among individuals with both phenotypes available; cross-region models used QTL representatives. Candidate interaction pairs comprised representatives within or between regions, requiring MAF ≥ 0.05, inter-locus *r*^2^ ≤ 0.2 and an estimable product effect. Sharing and interactions used the model, priors and 0.95 threshold described above.

### Combination ranking and allele-pyramiding design

Allele combinations were assessed from configuration and effect posteriors on the training-sample centring scale. For combinations *i* and *j* in the same genetic background, the genetic-effect difference for trait *t* is

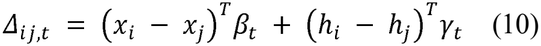

*x* and *h* are main-effect and interaction design vectors, with coefficients β_*t*_ and γ_*t*_. Users set improvement direction *d*_*t*_ to 1 for an increase or −1 for a decrease. We defined the posterior probability of simultaneous improvement (PPSI) of combination *i* relative to combination *j* as

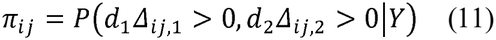

For *J* candidate combinations, the ranking score *S_i_* was defined as the mean PPSI of combination *i* relative to the other *J*-1 combinations:

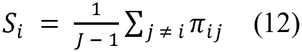

Posterior samples of genetic configurations and conditional effect distributions were used to estimate PPSI, mean PPSI and mean trait changes. Trait changes and their 95% equal-tailed posterior intervals were expressed in the original units. The double-low application used 201 Bna1007 accessions and ranked 16 homozygous combinations at four markers for reductions in EA and GSL. The other 161 candidate loci were fixed at reference states. C12, the most frequent observed combination, provided the effect-display reference. Complementary parents were identified based on cultivar provenance, four-marker genotypes and recorded EA and GSL phenotypes.

## Results

### An integrated framework for genetic analysis and breeding

We developed BPEA, an integrated framework for QTL interval identification, joint Bayesian inference of pleiotropy and epistasis, and allele-combination assessment (**Fig. 1**). Candidate intervals are defined using GWAS summary statistics and local LD structure, with boundary stability assessed by resampling individuals. Variants within these intervals and candidate interaction pairs are then analyzed in a bivariate Bayesian model that accounts for genetic relatedness and covariance between traits. The model jointly estimates main and epistatic effects while evaluating whether main-effect loci are shared between traits or trait-specific. Support for shared signals and candidate interactions is quantified by regional sharing probabilities and interaction posterior inclusion probabilities (PIPs), respectively.

**Figure 1.**
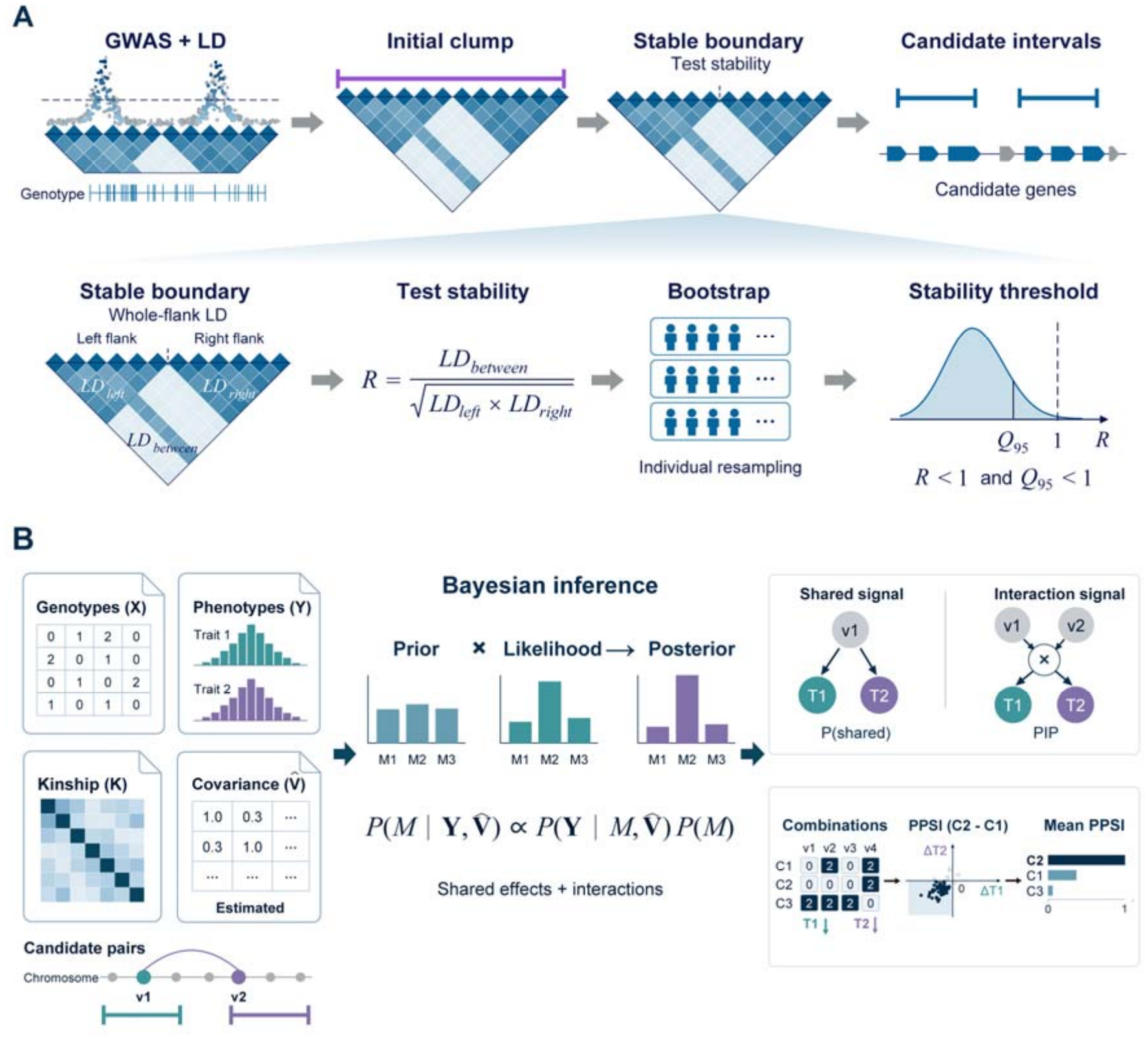
BPEA analysis framework. (A) QTL interval identification using genome-wide association studies (GWAS) and linkage disequilibrium (LD), with individual-level bootstrap assessment of boundary stability. *R* is mean cross-flank *r*^2^ divided by the geometric mean of within-flank mean *r*^2^; a boundary is retained when *R* and its bootstrap 95th percentile are below 1. (B) Inputs, model and outputs of joint Bayesian inference. M denotes the joint configuration of main and interaction effects; Y is the phenotype matrix and V□ the estimated covariance matrix. *P*(shared) is the regional posterior sharing probability; PIP is the posterior inclusion probability for a candidate interaction pair acting on a specified trait; PPSI denotes the posterior probability that one combination improves both target traits in the specified directions relative to another under the same genetic background. **Alt text:** Two-panel schematic illustrating the BPEA analysis framework. Panel A shows the identification of candidate QTL intervals from GWAS signals and local linkage disequilibrium, followed by bootstrap assessment of boundary stability. Panel B illustrates joint Bayesian analysis of candidate genetic variants and their interactions across two traits. The model accounts for genetic relatedness and trait covariance and produces posterior evidence for shared genetic signals and candidate epistatic interactions. Posterior genetic effects are further used to assess allele combinations for simultaneous improvement of target traits.

The joint posterior distribution also enables comparisons of allele combinations under the same genetic background. For each pair of allele combinations, BPEA estimates PPSI for simultaneous improvement of both traits in the specified directions. The model also reports mean trait changes and 95% posterior intervals. Candidate combinations are ranked by their mean PPSI relative to all other candidate combinations. Marker genotypes in existing accessions can then be used to identify parents carrying complementary target alleles. BPEA thus extends joint genetic inference to allele-combination assessment while propagating uncertainty in genetic configurations and effects.

### Interval identification in independent simulations

To evaluate QTL interval identification, we compared BPEA with p-clump and fixed-window approaches using the same GWAS summary statistics from 600 independently simulated genotype datasets (**Fig. 2; Supplementary Fig. 1; Supplementary Table 1–3**). The simulations encompassed six LD scenarios that varied in region length, internal LD heterogeneity and LD between neighbouring regions. Under default settings, BPEA achieved interval recall of 94.33% and 100% at phenotypic variance explained (PVE) levels of 8% and 16%, respectively, compared with 73.11% and 78.67% for p-clump. The corresponding precision values were 90.42% and 91.46% for BPEA and 75.11% and 68.67% for p-clump. BPEA therefore achieved higher recall and precision than p-clump at both PVE levels.

**Figure 2.**
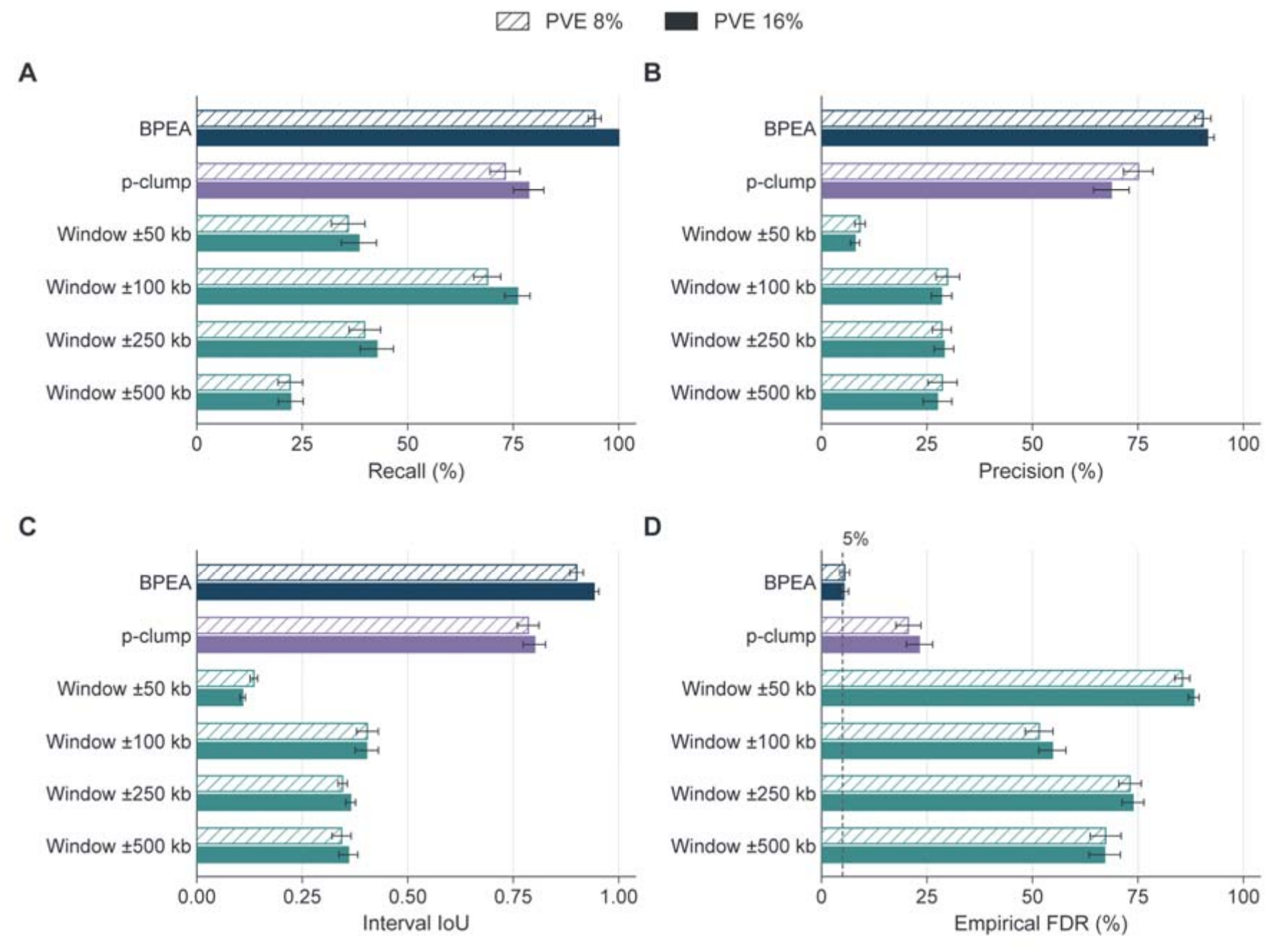
QTL interval identification in independent simulations. (A) Recall. (B) Precision. (C) Interval intersection over union (IoU), penalizing unmatched intervals. (D) Empirical interval false discovery rate (FDR), averaged across phenotype-specific false discovery proportions; the dashed line marks 5%. BPEA is compared with PLINK p-clump and fixed windows of ±50, ±100, ±250 and ±500 kb. The 600 independent genotype datasets each generated phenotypes at 8% and 16% phenotypic variance explained (PVE), with 900 target intervals at each level. LD uses Pearson *r*^2^ > 0.2. Recall and precision use one-to-one matching at IoU ≥ 0.5. Error bars indicate 95% confidence intervals from 10,000 paired bootstrap replicates. **Alt text:** Four-panel comparison of BPEA, PLINK p-clump and fixed-window approaches for identifying simulated QTL intervals. Panels A and B show interval recall and precision, respectively, with BPEA outperforming the comparator methods at both simulated effect sizes. Panel C shows greater agreement between BPEA intervals and target intervals, measured by intersection over union. Panel D shows lower empirical false discovery rates for BPEA than for p-clump. The comparisons summarize interval-identification performance across simulated linkage disequilibrium structures.

We further assessed agreement with simulated target intervals and false discoveries using the interval intersection-over-union (IoU) score and empirical interval false discovery rate (FDR). At 8% and 16% PVE, interval-IoU scores were 0.900 and 0.941 for BPEA, respectively, compared with 0.785 and 0.801 for p-clump. The mean paired improvement was 0.128 (95% bootstrap confidence interval, 0.107–0.149). The corresponding empirical interval FDRs were 5.47% and 5.31% for BPEA and 20.58% and 23.15% for p-clump, representing a mean reduction of 16.48 percentage points (95% bootstrap confidence interval, 13.97–19.05). Under the null, each method reported at least one interval in 29 of the 600 simulated phenotypes (4.83%; **Supplementary Fig. 2**). These results indicate that BPEA achieved greater agreement with simulated target intervals and lower empirical interval FDR than p-clump under the non-null conditions tested.

We also compared fixed windows extending 50, 100, 250 or 500 kb on either side of each lead marker. At 8% PVE, recall ranged from 22.11% to 68.89% and precision from 9.10% to 29.85%; at 16% PVE, the corresponding ranges were 22.22–76.00% and 7.94–29.02%. Increasing window size did not consistently improve interval matching, highlighting the limitations of fixed-window approaches when target intervals differ in length and local LD structure.

We next examined the sensitivity of interval identification to the LD threshold. Increasing the threshold to *r*^2^ > 0.4 reduced BPEA precision to 26.14% and 16.73% at 8% and 16% PVE, respectively. At *r*^2^ > 0.8, no reported interval met the IoU ≥ 0.5 matching criterion (**Supplementary Fig. 3**). These results indicate that more stringent LD thresholds do not necessarily improve interval identification.

**Figure 3.**
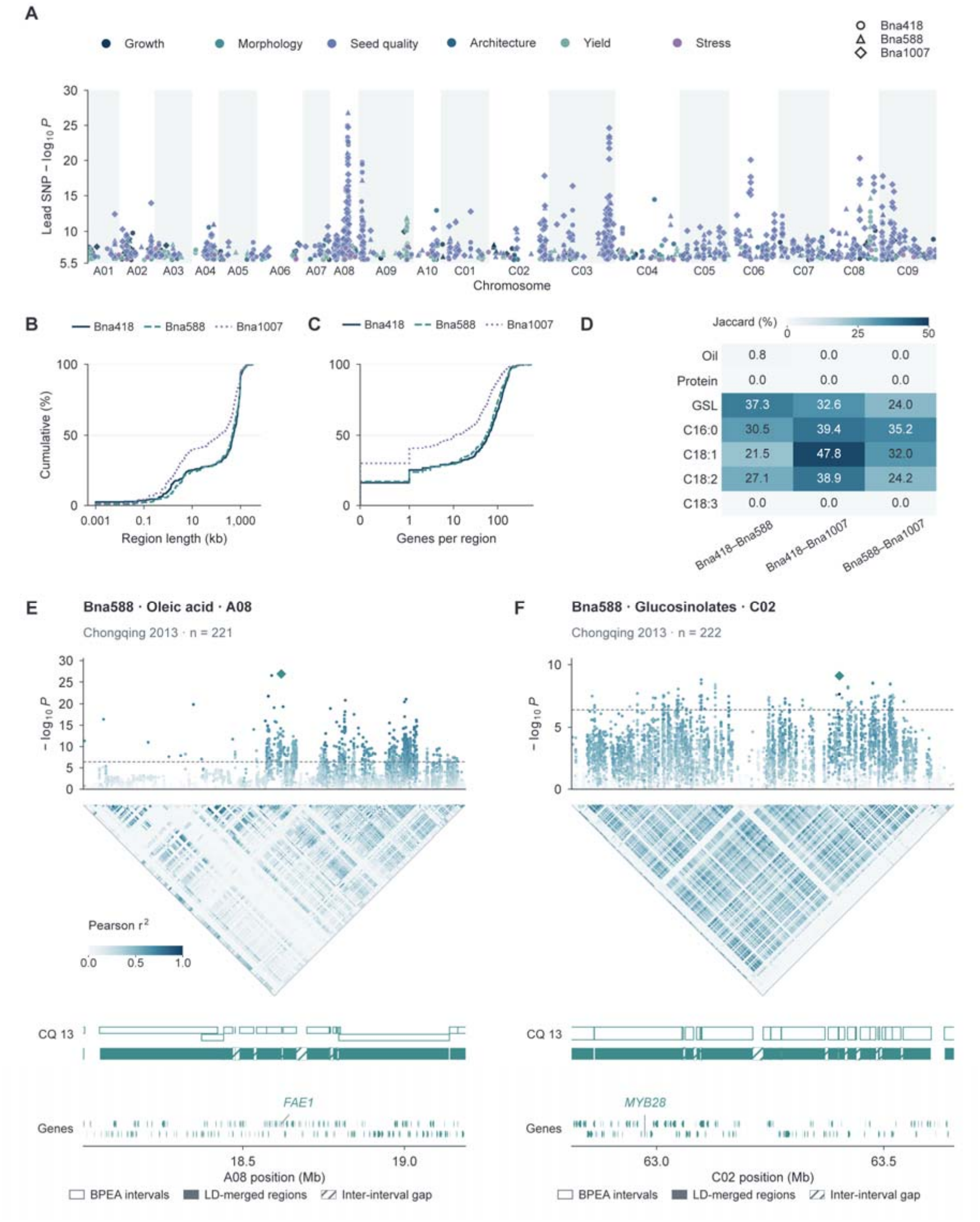
QTL atlas and regional cases in three rapeseed populations. (A) GWAS −log_10_*P* of QTL lead markers across 19 chromosomes; colours and shapes indicate trait classes and populations. (B) Cumulative distribution of region lengths. (C) Cumulative distribution of candidate-gene counts. (D) Cross-population Jaccard similarity of interval coverage for seven shared seed-quality traits. (E) Oleic-acid association signals and *FAE1* on A08 in Bna588 grown in Chongqing in 2013 (*n* = 221). (F) Glucosinolate association signals and *MYB28* on C02 in the same population and year (*n* = 222). Outlined bars, filled bars and hatching denote BPEA intervals, LD-merged regions and inter-interval gaps; arrows indicate gene orientation. Points in E and F are coloured by Pearson genotype *r*^2^ with the strongest associated marker in each window, shown as a diamond. Heatmaps show pairwise *r*^2^. Dashed lines mark *P* = 3.97 × 10□ □. **Alt text:** Six-panel overview of QTL intervals identified in three rapeseed populations. Panel A displays the genomic distribution of trait-associated signals across 19 chromosomes. Panels B and C show cumulative distributions of QTL interval lengths and positional candidate-gene counts, with Bna1007 generally exhibiting shorter intervals and fewer candidate genes. Panel D compares genomic interval overlap among populations for seven seed-quality traits. Panels E and F show regional association signals and linkage disequilibrium patterns surrounding *FAE1* and *MYB28*, respectively, illustrating the recovery of established seed-quality genes within candidate QTL intervals.

### A QTL and candidate-gene atlas across rapeseed populations

To further evaluate the applicability of BPEA’s interval-identification method to real populations, we identified and annotated candidate QTL intervals across 138 trait–environment combinations from three rapeseed populations, encompassing phenology, plant architecture, morphology, yield components, seed quality, and stress and disease responses. The resulting atlas comprised 1,358 QTL-region records, including 519 in Bna418, 371 in Bna588 and 468 in Bna1007 (**Fig. 3A; Supplementary Table 4**). Median region lengths were 506.783, 454.912 and 133.724 kb, respectively, with corresponding median counts of 63, 54 and 10.5 positional candidate genes per region (**Fig. 3B, C; Supplementary Table 5**). Bna1007 exhibited shorter candidate intervals and fewer positional candidate genes per region than the other two populations.

To assess the cross-population consistency of candidate QTL intervals, we compared their genomic coverage for seven seed-quality traits shared among the three rapeseed populations. Oleic acid showed the greatest overlap between Bna418 and Bna1007 (Jaccard index = 47.75%), whereas protein and linolenic acid showed no overlap in any of the three pairwise population comparisons (**Fig. 3D**). These results indicate that positional consistency varies among traits, with overlapping oleic-acid intervals providing candidates for further investigation.

To evaluate the biological relevance of the identified QTL intervals, we examined loci associated with oleic acid and glucosinolate content in Bna588 grown in Chongqing in 2013. The oleic acid-associated interval on A08 overlapped *FAE1*, which encodes an enzyme involved in very-long-chain fatty acid biosynthesis (James et al., 1995), with the lead marker located within the gene (*P* = 1.33 × 10□^2^□; *n* = 221; **Fig. 3E**). The glucosinolate-associated interval on C02 overlapped *MYB28*, a known regulator of seed glucosinolate accumulation (Zhou et al., 2023; *n* = 222; **Fig. 3F**). The original intervals identified in Bna418 and Bna1007 also overlapped these genes, including those associated with oleic acid (Yangluo 2014) and glucosinolates (Nanchang 2014) in Bna418, and erucic acid and glucosinolates (Zhejiang 2019) in Bna1007 (**Supplementary Fig. 4**). The recovery of these established seed-quality genes across populations supports the biological relevance of the identified candidate intervals.

**Figure 4.**
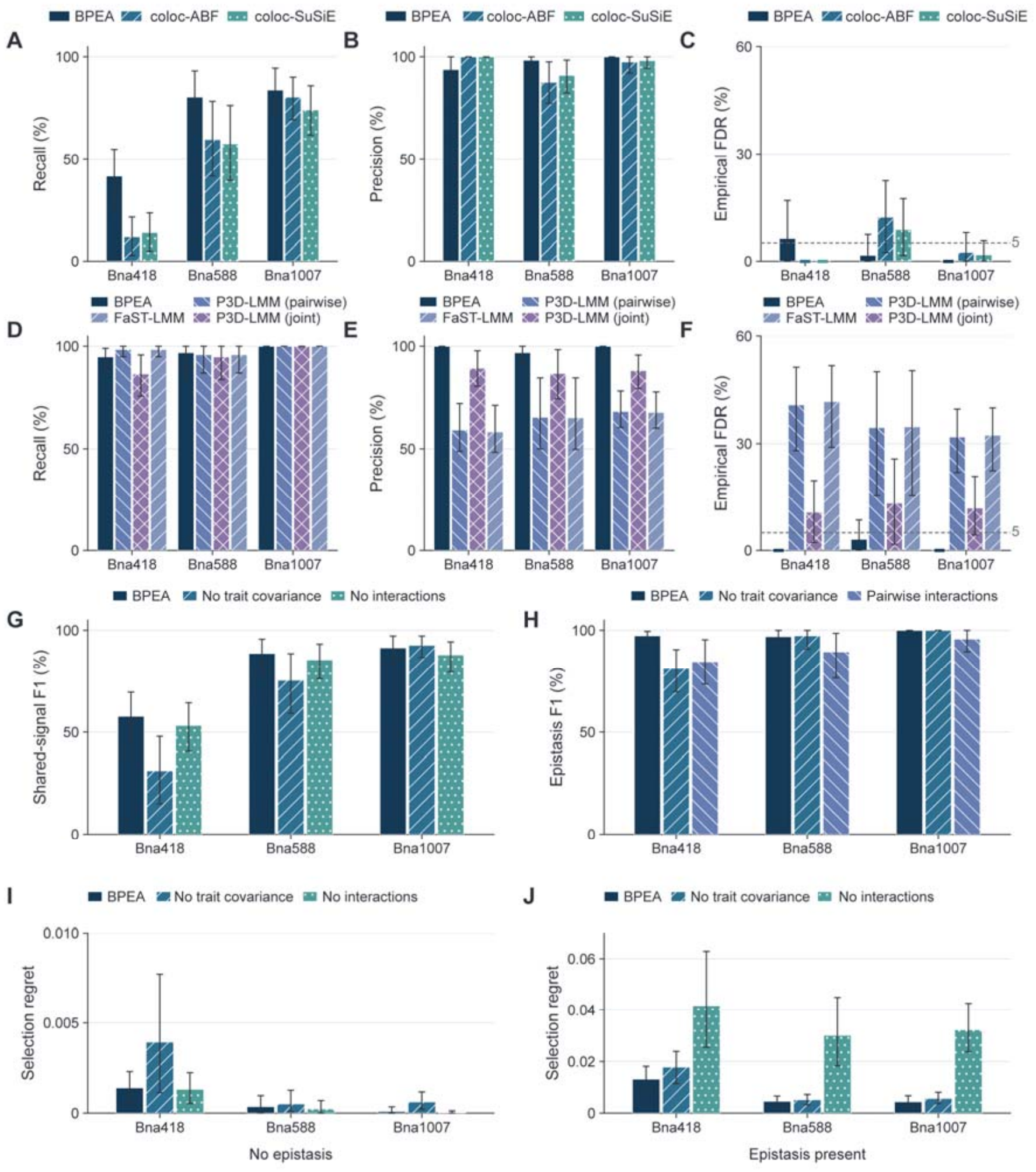
Joint-model accuracy, component contributions and combination selection. (A–C) Recall, precision and empirical FDR for shared-signal identification. (D–F) Recall, precision and empirical FDR for exact interaction-pair identification. (G) Shared-signal *F*_*1*_ in model ablations. (H) Interaction *F*_*1*_ in model ablations. (I) Selection regret without simulated epistasis. (J) Selection regret with simulated epistasis. Regret is the difference in mean true joint-improvement score between the optimal selection and the model-selected top 20%; lower is better. No trait covariance removes between-trait covariance; No interactions remove interaction terms; Pairwise interactions retain candidate main effects and fits one interaction pair at a time. A–H include 864 simulations across three populations; I and J include 863 diagnostic-eligible paired cases, each evaluating 64 observed genotype profiles. Error bars indicate 95% intervals from 2,000 paired region–template bootstrap resamples. Bayesian posterior thresholds in A–H are 0.95; interaction-test comparators use BH *q* ≤ 0.05. FDR dashed lines mark 5%. **Alt text:** Ten-panel evaluation of BPEA using simulations based on three rapeseed populations. Panels A–C compare recall, precision and empirical false discovery rate for shared genetic signal detection. Panels D–F evaluate the identification of specific epistatic interactions, showing that BPEA achieves higher precision than comparator methods while maintaining high recall. Panels G and H assess the effects of removing model components or fitting interactions separately. Panels I and J compare allele-combination selection regret in simulations without and with epistasis, respectively. When epistasis is present, the full BPEA model reduces selection regret relative to the model without interaction terms.

### Joint-model accuracy and component contributions

To evaluate the accuracy of shared-signal inference, we compared BPEA with coloc-ABF and coloc-SuSiE across 864 bivariate phenotype simulations based on real genotypes from three rapeseed populations (**Fig. 4A–C; Supplementary Fig. 5; Supplementary Table 6**). At a posterior probability threshold of 0.95, BPEA achieved precision of 93.75%, 98.31% and 100% and recall of 41.67%, 80.56% and 84.03% in Bna418, Bna588 and Bna1007, respectively. In Bna588, BPEA increased recall by 20.83 and 22.92 percentage points relative to coloc-ABF and coloc-SuSiE, respectively (95% paired bootstrap confidence intervals, 8.33–34.03 and 7.64–38.19 percentage points). In Bna1007, however, the improvement over coloc-ABF was smaller (3.47 percentage points; 95% confidence interval, −4.86 to 11.82). Overall, BPEA maintained high precision across populations, whereas recall and improvements relative to comparator methods varied, with the clearest gain observed in Bna588.

**Figure 5.**
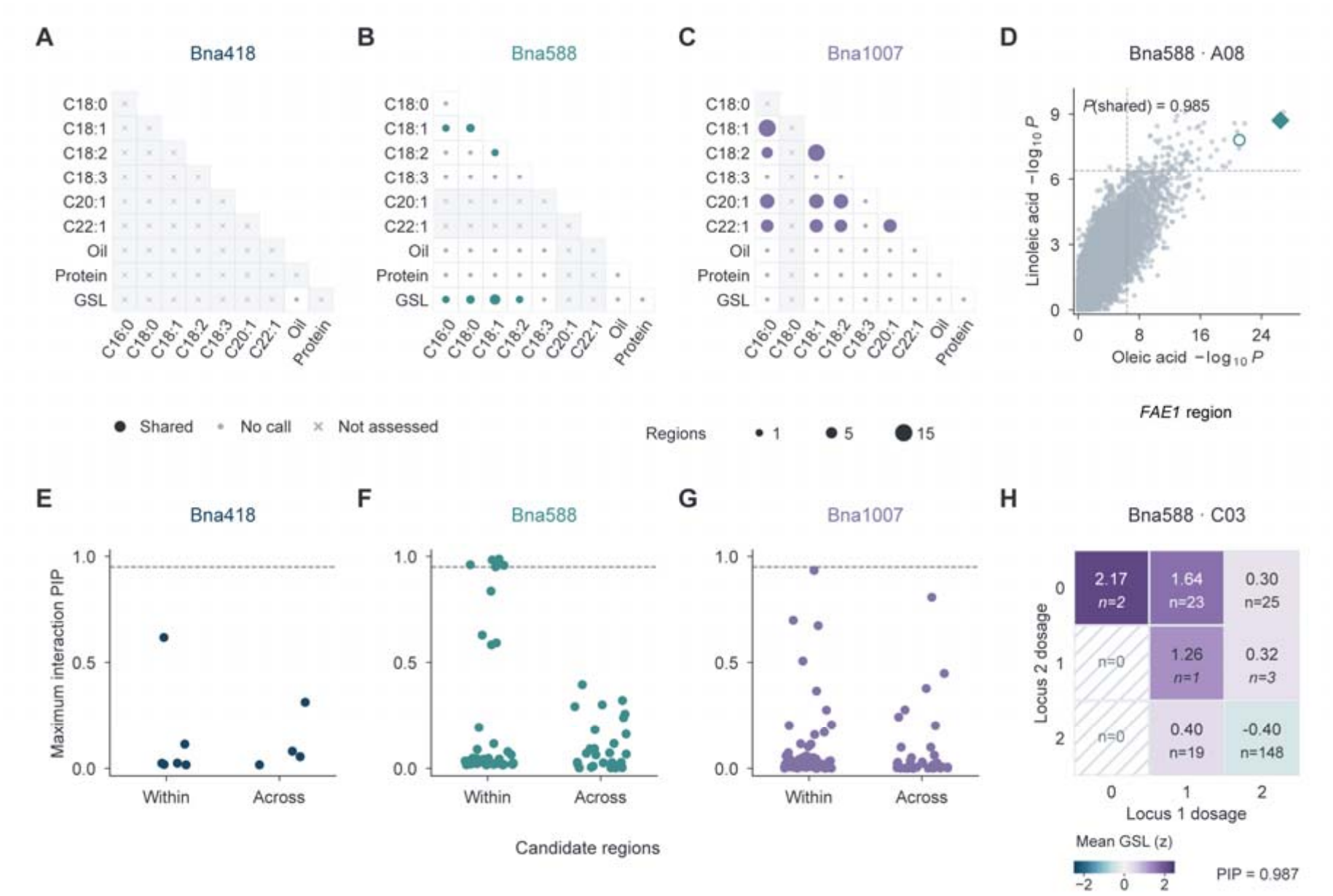
Shared genetic signals and epistasis for seed-quality traits. (A–C) Supported shared-region records in Bna418, Bna588 and Bna1007; circle area indicates record count, while grey dots and crosses denote no supported records and unevaluated pairs. (D) GWAS −log_10_*P* for oleic and linoleic acids at the same variants near *FAE1* on Bna588 A08, highlighting the two highest shared-variant PIPs; *n* = 221, regional *P*(shared) = 0.985, and dashed lines mark *P* = 3.97 × 10□□. (E–G) Maximum interaction PIP in within-region and cross-region models in the three populations. (H) The Bna588 C03:72,609,554 × 74,081,762 glucosinolate interaction (PIP = 0.987). Heatmap cells show observed standardized phenotype means and sample counts; hatching denotes no samples and italic *n* denotes *n* < 5. Posterior support thresholds for sharing and interactions are both 0.95. **Alt text:** Eight-panel visualization of shared genetic signals and candidate epistatic interactions associated with rapeseed seed-quality traits. Panels A–C summarize supported shared genetic signals across trait pairs in Bna418, Bna588 and Bna1007, with supported records detected in Bna588 and Bna1007 but not Bna418. Panel D shows association signals for oleic and linoleic acids near *FAE1* on chromosome A08, supporting a shared genetic association in this region. Panels E–G display posterior support for candidate epistatic interactions across the three populations. Panel H shows glucosinolate phenotypes across genotype combinations for a candidate interaction between two variants on chromosome C03 in Bna588.

To evaluate the identification of specific epistatic interactions, we compared BPEA with pairwise P3D-LMM, joint P3D-LMM and FaST-LMM using identical candidate sets (**Fig. 4D–F**). Across Bna418, Bna588 and Bna1007, BPEA achieved interaction recall of 94.79%, 96.88% and 100%, respectively, compared with 98.44%, 95.83% and 100% for FaST-LMM. The corresponding precision values were 100%, 96.88% and 100% for BPEA, substantially higher than 58.33%, 65.25% and 67.61% for FaST-LMM. Precision ranged from 59.25% to 68.09% for pairwise P3D-LMM and from 86.67% to 89.25% for joint P3D-LMM across the three populations. Overall, BPEA maintained high recall across all three populations while achieving higher precision and lower empirical FDR than all comparator methods, indicating fewer false-positive assignments of epistatic interactions to candidate variant pairs.

To evaluate the contributions of individual model components, we conducted ablation analyses across the same 864 simulated cases, removing trait covariance or interaction terms or fitting candidate interactions separately (**Fig. 4G, H; Supplementary Table 6**). Excluding trait covariance changed shared-signal recall by −22.92, −16.67 and +3.47 percentage points in Bna418, Bna588 and Bna1007, respectively. Removing interaction terms reduced the F_1_ score for shared-signal detection by 0.0305–0.0449 across the three populations, with 95% paired bootstrap confidence intervals for the full-model advantage excluding zero. Fitting candidate interactions separately preserved interaction recall but increased empirical FDR from 0%, 3.13% and 0% under the full model to 23.85%, 16.96% and 8.13% in Bna418, Bna588 and Bna1007, respectively. These results support the inclusion of epistatic effects in shared-signal inference and the joint evaluation of candidate interactions to reduce false-positive assignments, while indicating that the contribution of trait covariance varies across populations.

We next evaluated whether modeling epistasis improves allele-combination selection. In each simulation, 64 observed genotype profiles were ranked by mean PPSI for simultaneous reductions in both traits. The selections were assessed against their known simulated genetic effects. Across 863 simulations with paired results, selection regret was similar between BPEA and the model without interaction terms when epistasis was absent (**Fig. 4I**). In contrast, when epistasis was present, BPEA reduced mean selection regret from 0.04173 to 0.01318 in Bna418, from 0.03017 to 0.00459 in Bna588, and from 0.03231 to 0.00442 in Bna1007, corresponding to reductions of 68.4%, 84.8% and 86.3%, respectively (**Fig. 4J**). These results demonstrate that incorporating epistatic effects can improve allele-combination selection when interactions contribute to trait variation.

We also benchmarked the computational runtime of BPEA and the comparator methods under identical single-threaded conditions (**Supplementary Fig. 5C; Supplementary Table 7**). Median runtimes per case for BPEA were 5.022, 2.267 and 4.569 s in Bna418, Bna588 and Bna1007, respectively. Among the comparators, coloc-ABF and pairwise P3D-LMM were the fastest for shared-signal detection and epistasis testing, respectively. Their corresponding combined median runtimes per case were 1.019, 1.103 and 2.928 s. Overall, BPEA achieved higher precision in identifying epistatic interactions with a 1.16–4.00 s increase in median runtime per case compared with performing the two analyses separately.

### Pleiotropy and epistasis underlying seed quality

We next applied BPEA to 71 seed-quality trait pairs measured within the same population and environment to investigate shared genetic signals and epistatic interactions across three rapeseed populations. At a regional sharing probability threshold of 0.95, 106 region-by-trait-pair records met the criterion, including 94 records across 19 QTL regions in Bna1007 and 12 across six regions in Bna588; no records met the threshold in Bna418 (**Fig. 5A–C; Supplementary Table 8**). In Bna588, oleic and linoleic acid showed strong evidence of a shared genetic signal within an A08 interval (14,542,445–20,299,202 bp; regional sharing probability = 0.984875) based on 221 accessions with paired phenotypic data. Among the candidate variants, A08:18,589,228 had the highest shared posterior inclusion probability (PIP = 0.563875) and was located 28,824 bp from the annotated *FAE1* boundary, followed by A08:17,954,573 (shared PIP = 0.23525; **Fig. 5D**). Given the established role of *FAE1* in very-long-chain fatty-acid biosynthesis (James et al., 1995), these findings support a shared genetic association near *FAE1*.

We further investigated epistatic interactions affecting seed-quality traits in 192 joint models containing eligible candidate variant pairs, while accounting for main effects, genetic relatedness and trait covariance. At an interaction PIP threshold of 0.95, five records met the support criterion, all involving glucosinolate content and corresponding to two distinct variant pairs on chromosome C03 in Bna588 (**Fig. 5E–H; Supplementary Table 10**). The pair C03:72,609,554 x 73,301,346 had PIPs of 0.9505–0.9830 across four trait-pair models, whereas C03:72,609,554 x 74,081,762 had a PIP of 0.98725 in the oleic acid–glucosinolate model. Beyond seed-quality traits, we applied BPEA to plant architecture and yield components using 1,597 joint models. Nine interaction records met the posterior support threshold of 0.95, whereas no regional sharing records reached this threshold (**Supplementary Figs. 6 and 7; Supplementary Table 9 and 10**). These results indicate that candidate epistatic interactions can be detected in other agronomic traits even when evidence for shared genetic signals does not reach the same threshold.

**Figure 6.**
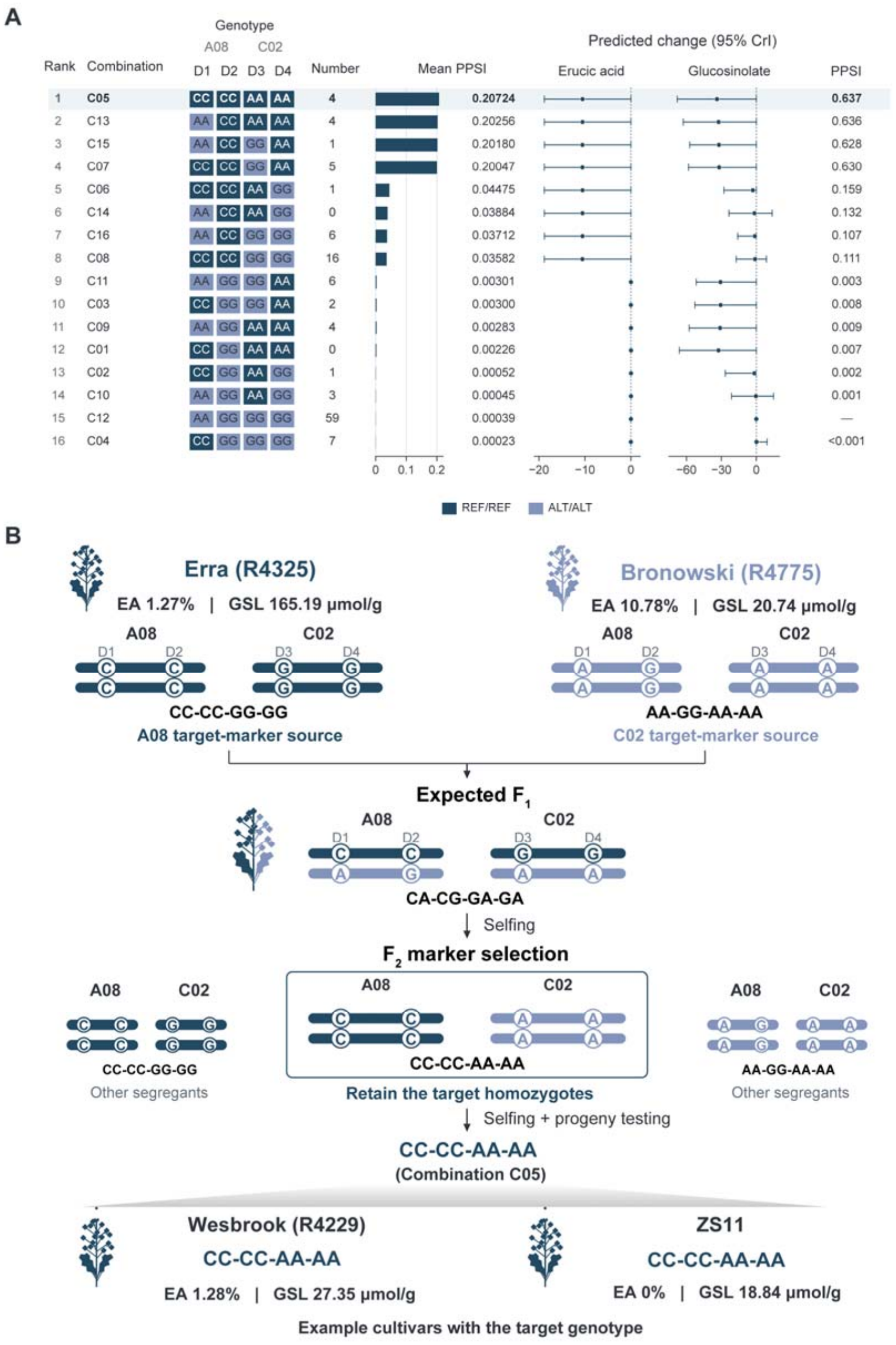
Combination ranking and a double-low allele-pyramiding design. (A) Ranking of 16 four-marker homozygous combinations. Scores are mean PPSI of jointly reducing erucic acid (EA) and glucosinolates (GSL) relative to the other 15 combinations; number denotes observed accession counts. Points and lines show posterior mean trait changes and 95% credible intervals relative to C12 (AA–GG–GG–GG), with the simultaneous-reduction probability shown separately. Other model loci are fixed at reference genotypes. (B) Proposed cross and marker-selection route between Erra (CC–CC–GG–GG) and Bronowski (AA–GG–AA–AA), targeting C05 (CC–CC–AA–AA). Wesbrook and ZS11 illustrate existing cultivars with the target marker genotype. **Alt text:** Two-panel illustration of allele-combination prioritization and proposed parental crossing for double-low rapeseed breeding. Panel A ranks 16 four-marker homozygous allele combinations by their mean posterior probability of simultaneous improvement (mean PPSI) for reducing erucic acid and glucosinolate contents. C05 (CC–CC–AA–AA) has the highest-ranking score. Predicted changes in both traits and their posterior intervals are shown relative to the reference combination C12. Panel B illustrates a proposed cross between Erra and Bronowski, which carry complementary target alleles, followed by selection of the C05 combination in segregating generations. Wesbrook and ZS11 are shown as existing cultivars carrying the target four-marker genotype.

### Allele pyramiding towards a double-low breeding target

To explore the application of BPEA in double-low rapeseed breeding, we used the joint posterior from Bna1007 to rank 16 homozygous four-marker allele combinations for simultaneous reductions in erucic acid (EA) and glucosinolate (GSL) contents. The C05 (CC–CC–AA–AA) combination ranked highest, with a mean PPSI of 0.20724, followed by C13, C15 and C07 (0.20256, 0.20180 and 0.20047, respectively; **Fig. 6A**).

To further characterize the predicted effects of the top-ranked allele combination, we compared C05 (CC–CC–AA–AA) with C12 (AA–GG–GG–GG), the most frequent observed four-marker combination, while holding the remaining candidate loci at their reference states. The PPSI of simultaneously reducing erucic acid (EA) and glucosinolate (GSL) contents was 0.63735. The model-averaged changes were −10.52 percentage points for EA and −33.80 μmol g□^1^ for GSL, indicating favorable predicted effects on both target traits.

To identify complementary parents for the prioritized C05 (CC–CC–AA–AA) allele combination, we examined cultivar provenance, four-marker genotypes and recorded EA and GSL phenotypes. Erra (R4325), carrying CC–CC–GG–GG, had EA and GSL values of 1.27% and 165.19 μmol g□^1^, respectively, whereas Bronowski (R4775), carrying AA–GG–AA–AA, had corresponding values of 10.78% and 20.74 μmol g□^1^. Erra and Bronowski carry complementary target alleles at the A08 and C02 loci, respectively (**Fig. 6B**). Previous studies have characterized Erra as a German winter-type cultivar and Bronowski as a Polish spring-type cultivar with low glucosinolate content (Fritsche et al., 2012; Josefsson and Appelqvist, 1968). A cross between these parents is expected to produce CA–CG–GA–GA F□ plants, from which the target C05 combination can be selected and fixed in segregating generations.

The prioritized C05 combination was also identified in existing cultivars with favorable seed-quality profiles. In Bna1007, Wesbrook (R4229) carried C05 and had recorded EA and GSL values of 1.28% and 27.35 μmol g□^1^, respectively. The same four-marker combination matched the ZS11 reference genotype, for which an erucic acid content of 0% and a glucosinolate content of 18.84 μmol g□^1^ were reported in national cultivar trials (Wang et al., 2009). Together, these observations provide additional support for the relevance of C05 to double-low rapeseed breeding.

## Discussion

The breeding value of an allele depends on the improvement objective and its genetic background. In tomato, negative epistasis between individually favorable alleles can reduce fertility (Soyk et al., 2017). Pleiotropic effects can also introduce trade-offs among traits, further complicating the selection of favorable allele combinations (Song et al., 2022). BPEA integrates LD-guided QTL interval identification with joint Bayesian inference of pleiotropy and epistasis, extending genetic dissection to allele-combination assessment while propagating uncertainty in genetic configurations and effects. Its application to rapeseed enabled the prioritization of allele combinations and complementary parents for reducing erucic acid and glucosinolate contents, two established seed-quality breeding targets (Wang et al., 2009).

Fixed windows and local LD blocks are commonly used to define candidate regions in crop GWAS (Knoch et al., 2020), but physical proximity or LD with a lead marker does not necessarily capture heterogeneity in local LD structure. BPEA addresses this limitation by evaluating LD differences across candidate boundaries and assessing their stability through bootstrap resampling. In simulations, BPEA achieved higher interval recall and precision than both p-clump and fixed-window approaches, along with greater agreement with target intervals and lower empirical FDR than p-clump. Additionally, increasing fixed-window size did not consistently improve interval matching, while sensitivity analyses showed that more stringent LD thresholds did not necessarily improve interval identification. Beyond statistical performance, interval delineation also has implications for identifying regulatory variants outside gene bodies. For example, a transposable element upstream of *BnaA9*.*CYP78A9* enhances gene expression and affects silique length and seed weight in rapeseed (Shi et al., 2019). The recovery of *FAE1* and *MYB28* in seed-quality QTL intervals across all three rapeseed populations further supports the biological relevance of the identified candidate regions.

Existing methods address different aspects of complex genetic inference. coloc-SuSiE accommodates multiple association signals during colocalization analysis (Wallace, 2021), whereas mvMAPIT uses trait correlations to improve the detection of marginal epistasis (Stamp et al., 2023). In BPEA, shared genetic signals and candidate epistatic interactions are evaluated within the same Bayesian model. Our ablation analyses showed that fitting candidate interactions separately increased empirical FDR without improving recall, while removing interaction terms reduced the F_1_ score for shared-signal detection. The contribution of trait covariance to shared-signal detection varied across datasets. These findings support joint inference of main and epistatic effects and provide a basis for propagating their posterior uncertainty into allele-combination assessment.

Recent genomic and breeding studies have highlighted the value of pyramiding favorable alleles for crop improvement (Gu et al., 2023; Yao et al., 2026). Genomics-guided breeding also requires suitable parental combinations and field evaluation of the resulting materials (Abdullah et al., 2026). BPEA supports combination selection by jointly evaluating genetic effects across traits and interactions between loci. In simulations with epistasis, BPEA reduced selection regret relative to a model without interaction terms, supporting the importance of modeling epistatic effects during combination selection. Rankings based on mean PPSI integrate genetic effects across traits. PPSI relative to a reference combination and predicted trait changes provide complementary information on the probability and magnitude of improvement. In rapeseed, BPEA prioritized C05 (CC–CC–AA–AA) for reducing erucic acid and glucosinolate contents and identified Erra and Bronowski as complementary parents carrying the target alleles. The presence of the same four-marker combination in Wesbrook and the ZS11 reference genotype, together with their favorable seed-quality profiles, provided further support for this breeding target. These findings demonstrate how joint genetic inference can inform parent selection and allele-pyramiding strategies.

Further development of BPEA could extend its application to more complex genetic architectures. The current candidate-selection strategy relies primarily on QTL-associated variants, potentially missing epistatic interactions involving loci without detectable marginal effects, as previously reported in rapeseed (Radoev et al., 2008). Expanding the search for candidate interactions and allowing multiple active interaction pairs within a genetic configuration, together with sensitivity analyses of prior specifications and covariance estimates, could broaden the scope of joint inference. For breeding applications, the prioritized C05 combination and the proposed Erra × Bronowski cross provide a basis for experimental validation. Tracking the segregation and fixation of target alleles, followed by comparisons with parental and control genotypes across environments, will help determine whether the predicted improvements translate into stable breeding gains.

## Author contributions

K.L. and X.L. conceived and designed the study. X.L. developed the methodology, performed the computational analyses and wrote the manuscript. Y.W., B.M., W.C., J.X. and Z.Q. curated the data. J.S. provided methodological guidance. K.L. supervised the research and acquired funding. J.S. and K.L. revised the manuscript. All authors reviewed and approved the final manuscript.

## Acknowledgements

We thank members of the Lu lab for their helpful feedback and discussion.

## Funding

This work was supported by the National Natural Science Foundation of China (32672682 and 32272111), the Chongqing Enterprise and Research Joint Commercial Breeding Innovation Project (CQZY-QKLH2026003), Chongqing Natural Science Foundation (CSTB2024NSCQ-LZX0012), Chongqing Modern Agricultural Industry-Technology System (CQMAITS202504), the 2025 Joint Construction Project of the Fuling Research Institute of Southwest University (FLYJY202504), Modern Agro-industry Technology Research System (CARS-12), and the National Key Research and Development Program of China (2022YFD1200400).

## Conflicts of interest

The authors declare no competing interests.

## Data availability statement

The public resequencing data are available from NCBI under BioProjects PRJNA476657 (Bna1007), PRJNA358784 (Bna588) and PRJNA416679 (Bna418). BPEA is available through GitHub at https://github.com/swu1019lab/BPEA.

## Notes

### Competing Interest Statement

The authors have declared no competing interest.

## References

Abdullah, M., Manzoor, M. A., Anwar, S., Henry, R. J. and Periyannan, S. (2026). From genes to farms: a modern breeding revolution. Trends in Plant Science. 10.1016/j.tplants.2026.09.012.

Benjamini, Y. and Hochberg, Y. (1995). Controlling the false discovery rate: a practical and powerful approach to multiple testing. Journal of the Royal Statistical Society: Series B (Methodological), 57, 289–300. 10.1111/j.2517-6161.1995.tb02031.x

Chang, C. C., Chow, C. C., Tellier, L. C. A. M., Vattikuti, S., Purcell, S. M. and Lee, J. J. (2015). Second-generation PLINK: rising to the challenge of larger and richer datasets. GigaScience, 4, 7. 10.1186/s13742-015-0047-8

Da Costa e Silva, L., Wang, S. and Zeng, Z.-B. (2012). Multiple trait multiple interval mapping of quantitative trait loci from inbred line crosses. BMC Genetics, 13, 67. 10.1186/1471-2156-13-67

Fritsche, S., Wang, X., Li, J. et al. (2012). A candidate gene-based association study of tocopherol content and composition in rapeseed (Brassica napus). Frontiers in Plant Science, 3, 129. 10.3389/fpls.2012.00129

Giambartolomei, C., Vukcevic, D., Schadt, E. E., Franke, L., Hingorani, A. D., Wallace, C. and Plagnol, V. (2014). Bayesian test for colocalisation between pairs of genetic association studies using summary statistics. PLOS Genetics, 10, e1004383. 10.1371/journal.pgen.1004383

Gu, Z., Gong, J., Zhu, Z. et al. (2023). Structure and function of rice hybrid genomes reveal genetic basis and optimal performance of heterosis. Nature Genetics, 55, 1745–1756. 10.1038/s41588-023-01495-8

Hu, J., Chen, B., Zhao, J. et al. (2022). Genomic selection and genetic architecture of agronomic traits during modern rapeseed breeding. Nature Genetics, 54, 694–704. 10.1038/s41588-022-01055-6

James, D. W., Jr., Lim, E., Keller, J., Plooy, I., Ralston, E. and Dooner, H. K. (1995). Directed tagging of the Arabidopsis FATTY ACID ELONGATION1 (FAE1) gene with the maize transposon activator. The Plant Cell, 7, 309–319. 10.1105/tpc.7.3.309

Josefsson, E. and Appelqvist, L.-Å. (1968). Glucosinolates in seed of rape and turnip rape as affected by variety and environment. Journal of the Science of Food and Agriculture, 19, 564–570. 10.1002/jsfa.2740191004

Knoch, D., Abbadi, A., Grandke, F. et al. (2020). Strong temporal dynamics of QTL action on plant growth progression revealed through high-throughput phenotyping in canola. Plant Biotechnology Journal, 18, 68–82. 10.1111/pbi.13171

Li, M. X., Yeung, J. M. Y., Cherny, S. S. and Sham, P. C. (2012). Evaluating the effective numbers of independent tests and significant p-value thresholds in commercial genotyping arrays and public imputation reference datasets. Human Genetics, 131, 747–756. 10.1007/s00439-011-1118-2

Lippert, C., Listgarten, J., Davidson, R. I., Baxter, J., Poon, H., Kadie, C. M. and Heckerman, D. (2013). An exhaustive epistatic SNP association analysis on expanded Wellcome Trust data. Scientific Reports, 3, 1099. 10.1038/srep01099

Liu, J., Hua, W., Hu, Z. et al. (2015). Natural variation in ARF18 gene simultaneously affects seed weight and silique length in polyploid rapeseed. Proceedings of the National Academy of Sciences of the USA, 112, E5123–E5132. 10.1073/pnas.1502160112

Mackay, T. F. C. and Anholt, R. R. H. (2024). Pleiotropy, epistasis and the genetic architecture of quantitative traits. Nature Reviews Genetics, 25, 639–657. 10.1038/s41576-024-00711-3

Qi, G., Li, Y., Zhang, W. et al. (2025). Reveal genomic insights into cotton domestication and improvement using gene level functional haplotype-based GWAS. Nature Communications, 16, 4734. 10.1038/s41467-025-59983-w

Radoev, M., Becker, H. C. and Ecke, W. (2008). Genetic analysis of heterosis for yield and yield components in rapeseed (Brassica napus L.) by quantitative trait locus mapping. Genetics, 179, 1547–1558. 10.1534/genetics.108.089680

Shi, L., Song, J., Guo, C. et al. (2019). A CACTA-like transposable element in the upstream region of BnaA9.CYP78A9 acts as an enhancer to increase silique length and seed weight in rapeseed. The Plant Journal, 98, 524–539. 10.1111/tpj.14236

Song, J.-M., Guan, Z., Hu, J. et al. (2020). Eight high-quality genomes reveal pan-genome architecture and ecotype differentiation of Brassica napus. Nature Plants, 6, 34–45. 10.1038/s41477-019-0577-7

Song, X., Meng, X., Guo, H. et al. (2022). Targeting a gene regulatory element enhances rice grain yield by decoupling panicle number and size. Nature Biotechnology, 40, 1403–1411. 10.1038/s41587-022-01281-7

Soyk, S., Lemmon, Z. H., Oved, M. et al. (2017). Bypassing negative epistasis on yield in tomato imposed by a domestication gene. Cell, 169, 1142–1155.e12. 10.1016/j.cell.2017.04.032

Stamp, J., DenAdel, A., Weinreich, D. and Crawford, L. (2023). Leveraging the genetic correlation between traits improves the detection of epistasis in genome-wide association studies. G3: Genes, Genomes, Genetics, 13, jkad118. 10.1093/g3journal/jkad118

Wallace, C. (2021). A more accurate method for colocalisation analysis allowing for multiple causal variants. PLOS Genetics, 17, e1009440. 10.1371/journal.pgen.1009440

Wang, G., Sarkar, A., Carbonetto, P. and Stephens, M. (2020). A simple new approach to variable selection in regression, with application to genetic fine mapping. Journal of the Royal Statistical Society: Series B (Statistical Methodology), 82, 1273–1300. 10.1111/rssb.12388

Wang, H.-Z., Liu, G.-H., Wang, X.-F., Liu, J., Yang, Q. and Hua, W. (2009). Heterosis and breeding of high oil content in rapeseed (Brassica napus L.). Proceedings of the 16th Australian Research Assembly on Brassicas, Ballarat, Victoria, Australia.

Wang, Y., Sun, D., Duan, Y. et al. (2025). A variome–transcriptome–metabolome network links GABA biosynthesis to stress resilience in maize. The Plant Cell, 37, koaf221. 10.1093/plcell/koaf221

Yao, W., Ku, L., Wang, B. et al. (2026). Breeding ideotype maize with enhanced yield through genomics-guided pyramiding of favorable alleles. Nature Genetics, 58, 916–927. 10.1038/s41588-026-02522-0

Zhang, Z., Ersoz, E., Lai, C. Q. et al. (2010). Mixed linear model approach adapted for genome-wide association studies. Nature Genetics, 42, 355–360. 10.1038/ng.546

Zhou, X. and Stephens, M. (2012). Genome-wide efficient mixed-model analysis for association studies. Nature Genetics, 44, 821–824. 10.1038/ng.2310

Zhou, X., Zhang, H., Xie, Z. et al. (2023). Natural variation and artificial selection at the BnaC2.MYB28 locus modulate Brassica napus seed glucosinolate. Plant Physiology, 191, 352–368. 10.1093/plphys/kiac463

