## Supplemental figures for "BPEA integrates pleiotropy and epistasis to guide allele pyramiding for multi-trait crop improvement"

### Supplementary Figure 1 | LD layouts and QTL interval identification

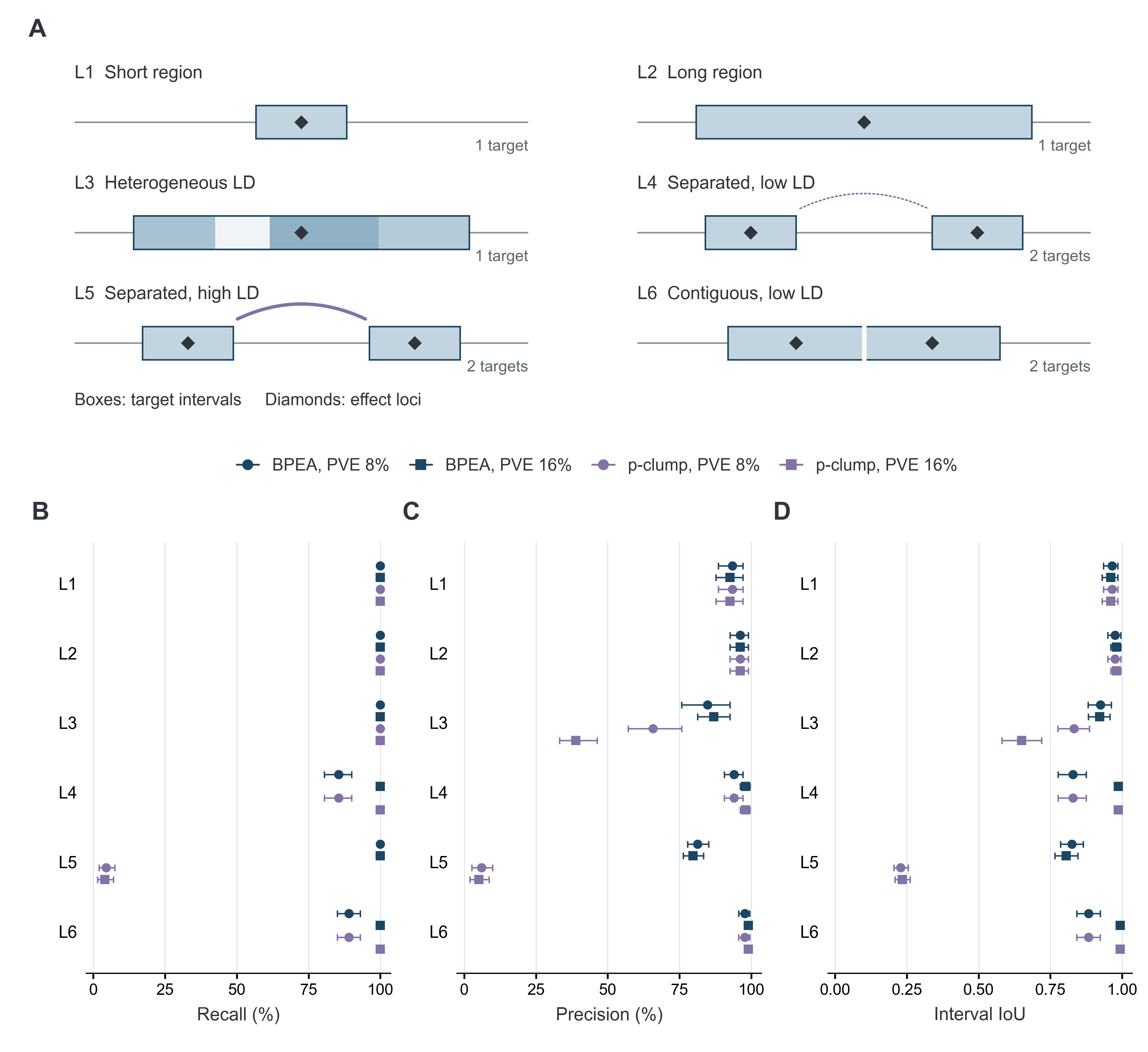

(A) Six LD layouts; boxes and diamonds denote target intervals and effect loci. (B) Recall. (C) Precision. (D) Interval IoU. Panels (B)–(D) show results at 8% and 16% PVE. Each layout contains 100 independent datasets. Points and error bars show estimates and 95% intervals from 10,000 paired bootstrap resamples within each layout. Default LD threshold: *r*² > 0.2; interval matching: IoU ≥ 0.5.

### Supplementary Figure 2 | False interval reports under the null

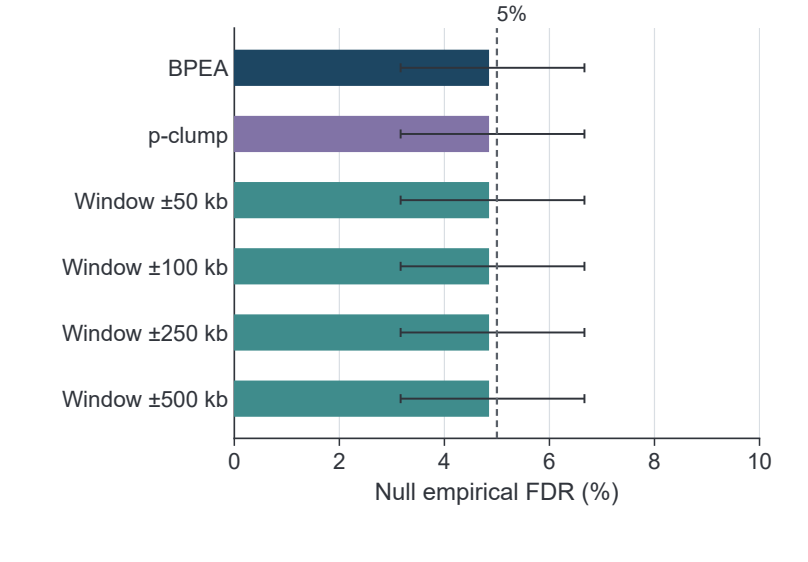

Bars show the proportion of 600 null phenotypes with at least one reported interval; error bars are 95% confidence intervals from 10,000 bootstrap replicates. The dashed line marks 5%.

### Supplementary Figure 3 | Effects of LD threshold and calculation mode on interval identification

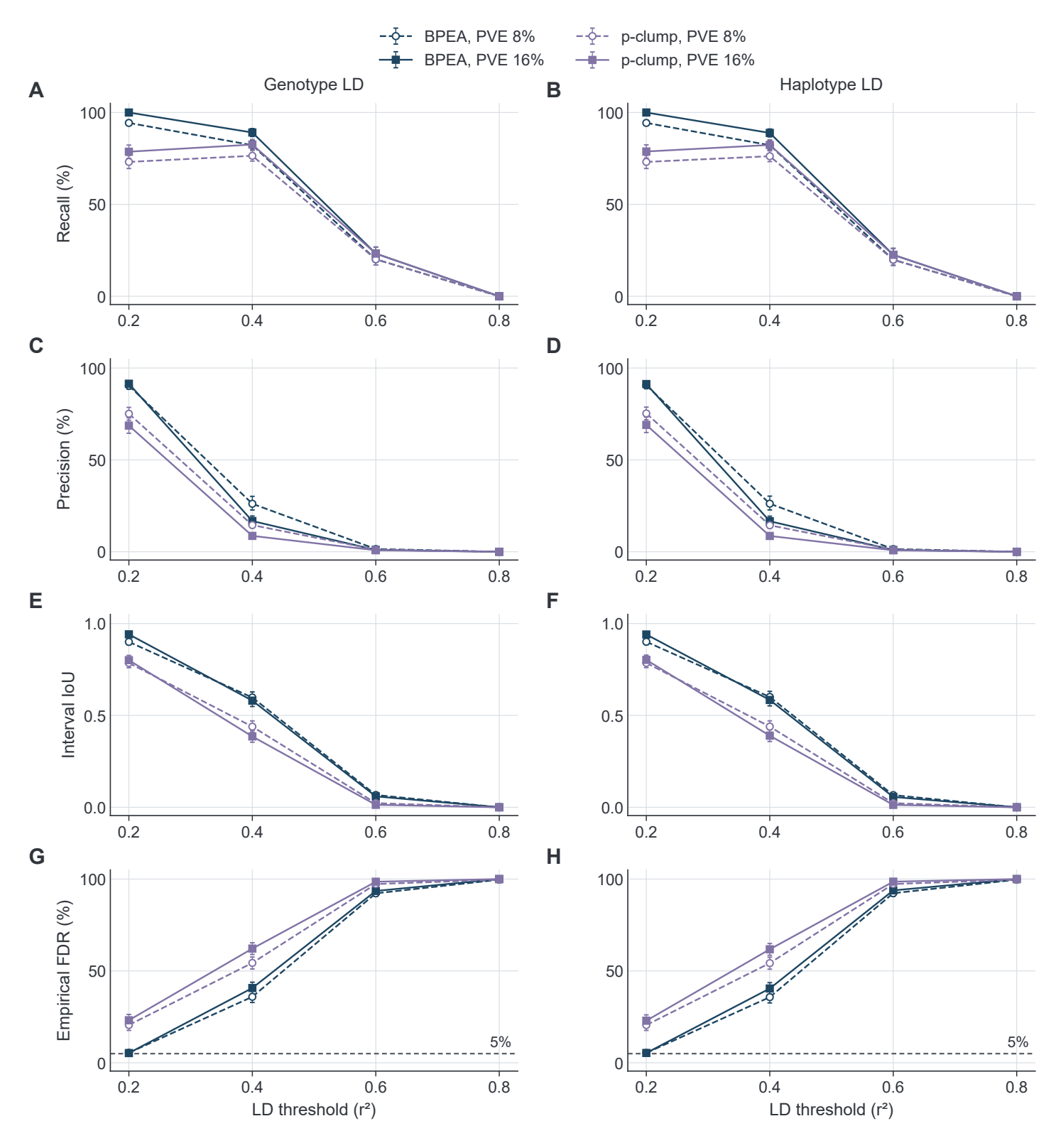

(A) Recall using genotype LD. (B) Recall using haplotype LD. (C) Precision using genotype LD. (D) Precision using haplotype LD. (E) Interval IoU using genotype LD. (F) Interval IoU using haplotype LD. (G) Empirical FDR using genotype LD. (H) Empirical FDR using haplotype LD. Genotype LD is based on allele-dosage correlations. Results are shown at 8% and 16% phenotypic variance explained (PVE). Error bars are 95% confidence intervals from 10,000 paired bootstrap replicates. Dashed FDR lines mark 5%.

### Supplementary Figure 4 | Quality-trait intervals overlapping *FAE1* and *MYB28* in Bna418 and Bna1007

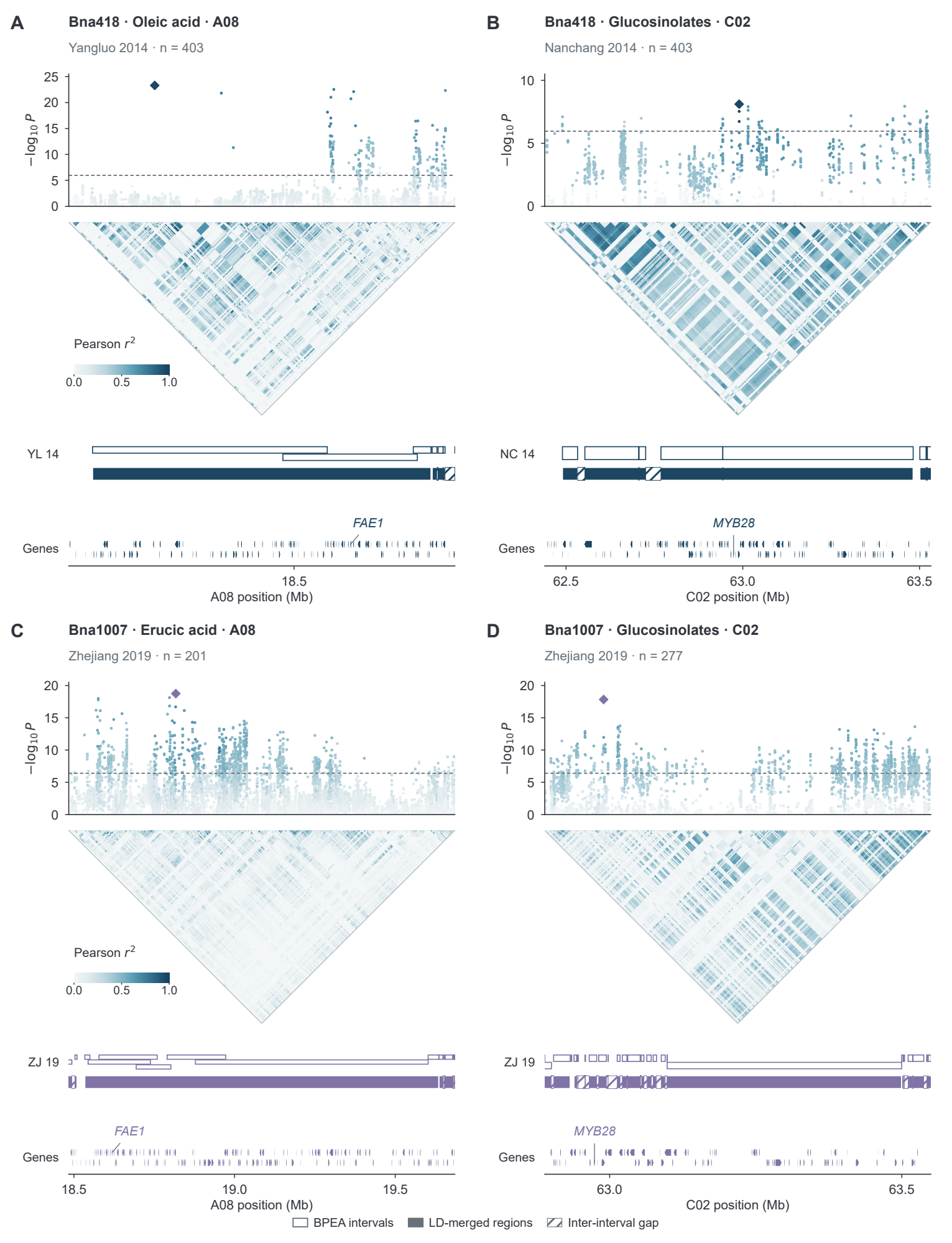

(A) Bna418 oleic acid–*FAE1*, Yangluo 2014 (*n* = 403). (B) Bna418 glucosinolates–*MYB28*, Nanchang 2014 (*n* = 403). (C) Bna1007 erucic acid–*FAE1*, Zhejiang 2019 (*n* = 201). (D) Bna1007 glucosinolates–*MYB28*, Zhejiang 2019 (*n* = 277). Symbols and interval tracks follow Fig. 3E,F. Dashed GWAS thresholds are *P* = 1.09 × 10⁻⁶ for Bna418 and *P* = 3.96 × 10⁻⁷ for Bna1007.

### Supplementary Figure 5 | Shared-signal architectures and computational runtime

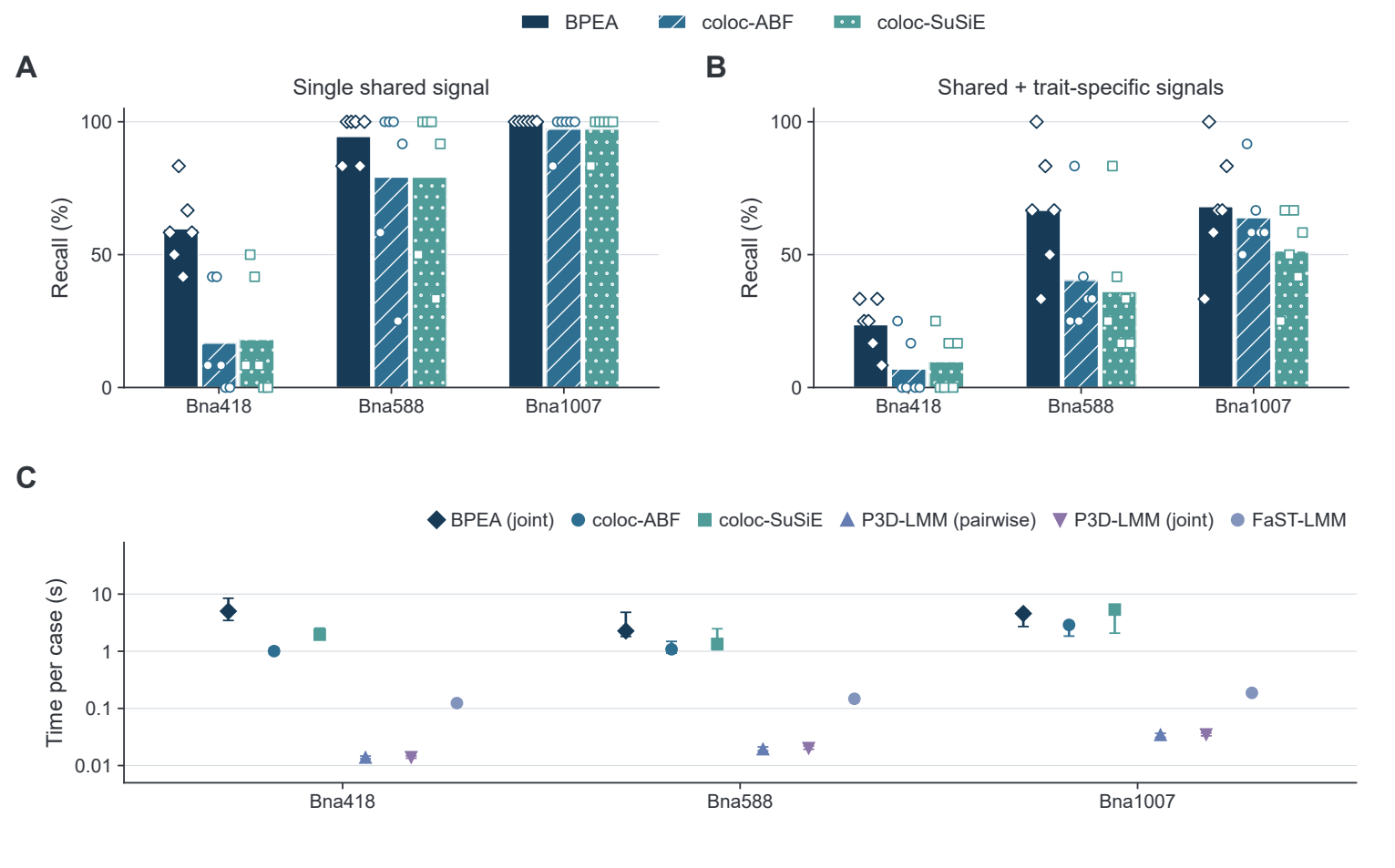

(A) A single shared signal. (B) A shared signal accompanied by a trait-specific signal. (C) Single-threaded per-case runtime, shown as median and interquartile range; BPEA timing includes both sharing and interaction inference. Bars and points in A and B show pooled and region-specific recall, respectively, at a posterior threshold of 0.95.

### Supplementary Figure 6 | Joint genetic analysis of additional trait classes

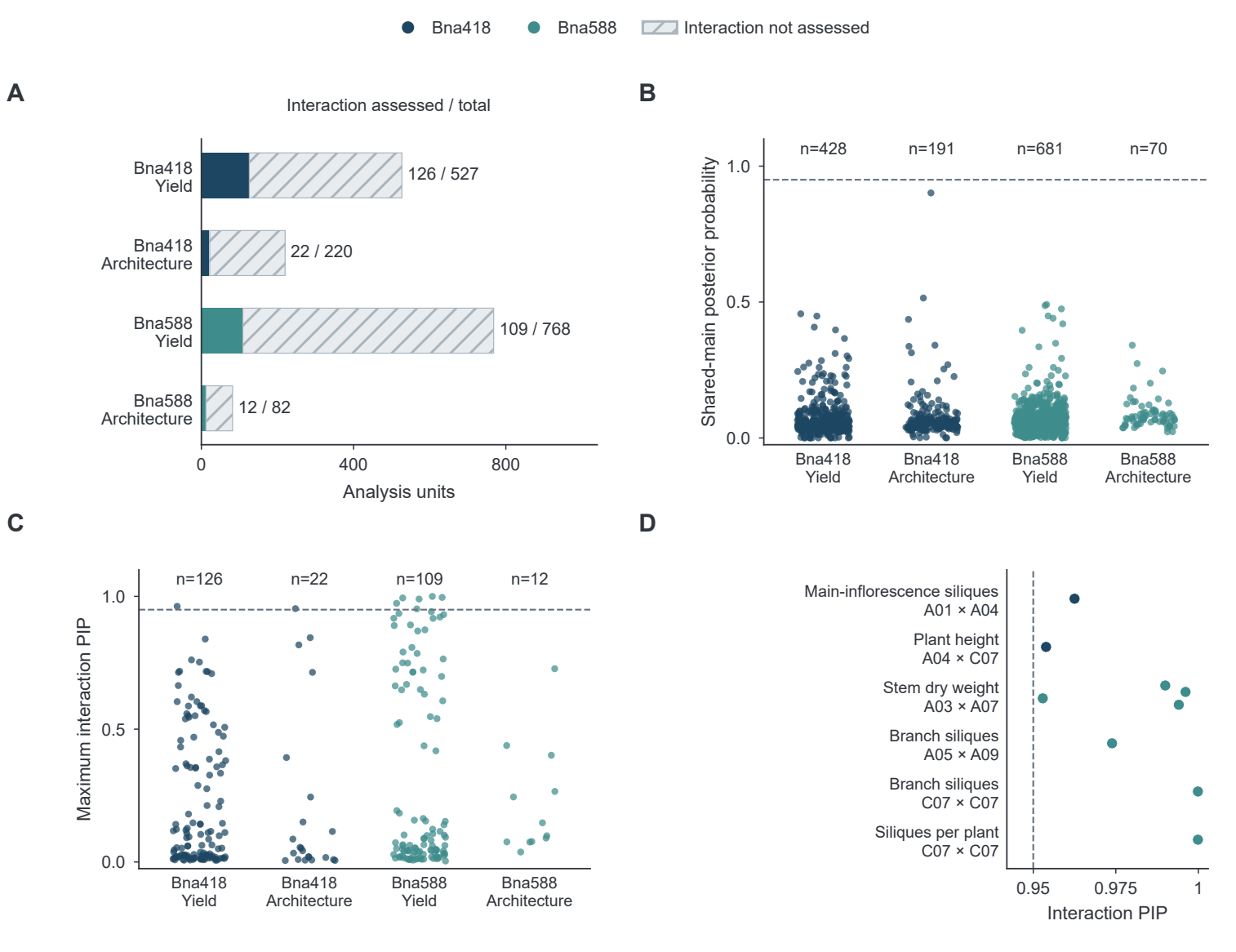

(A) Analysis coverage. (B) Regional sharing probabilities. (C) Maximum interaction posterior inclusion probability (PIP). (D) Supported interaction PIPs. Dashed lines indicate the 0.95 posterior threshold.

### Supplementary Figure 7 | Epistatic interactions and genotype phenotypes in additional traits

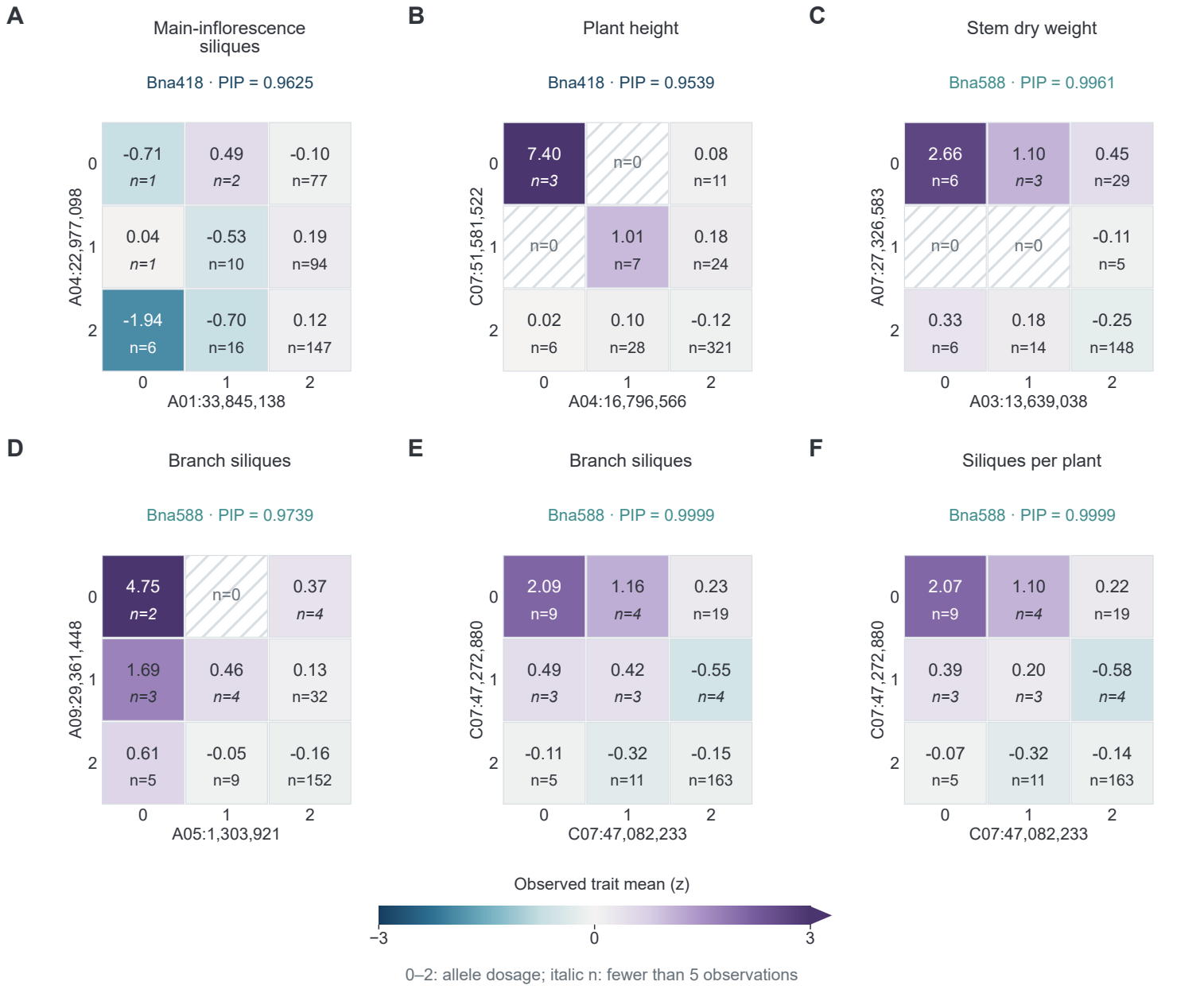

(A) Main-inflorescence siliques in Bna418. (B) Plant height in Bna418. (C) Stem dry weight in Bna588. (D) Branch siliques in Bna588. (E) Branch siliques in Bna588. (F) Siliques per plant in Bna588. Heatmaps show observed standardized phenotype means and sample counts for two-locus genotype combinations. Axes use allele dosages 0, 1 and 2; hatching indicates no observations and italic *n* indicates fewer than five observations. PIP is the posterior inclusion probability for the indicated trait; support requires PIP ≥ 0.95.
